# Extending the genotype in *Brachypodium* by including DNA methylation reveals a joint contribution with genetics on adaptive traits

**DOI:** 10.1101/840744

**Authors:** SR Eichten, A Srivastava, A Reddiex, DR Ganguly, A Heussler, J Streich, P Wilson, JO Borevitz

## Abstract

Epigenomic changes have been considered a potential missing link underlying phenotypic variation in quantitative traits but is potentially confounded with the underlying DNA sequence variation. Although the concept of epigenetic inheritance has been discussed in depth, there have been few studies attempting to directly dissect the amount of epigenomic variation within inbred natural populations while also accounting for genetic diversity. By using known genetic relationships between *Brachypodium* lines, multiple sets of nearly identical accession families were selected for phenotypic studies and DNA methylome profiling to investigate the dual role of (epi)genetics under simulated natural seasonal climate conditions. Despite reduced genetic diversity, appreciable phenotypic variation was still observable in the measured traits (height, leaf width and length, tiller count, flowering time, ear count) between as well as within the inbred accessions. However, with reduced genetic diversity there was diminished variation in DNA methylation within families. Mixed-effects linear modelling revealed large genetic differences between families and a minor contribution of epigenomic variation on phenotypic variation in select traits. Taken together, this analysis suggests a limited but significant contribution of DNA methylation towards heritable phenotypic variation relative to genetic differences.

## Introduction

Heritable natural variation has largely been attributed to genetic variation between individuals within and across populations. Novel combinations of alleles and regulatory sequences can influence gene expression and lead to complex changes in downstream phenotypes. Tools such as genome-wide association studies have been highly successful for identifying regions of the genome that contribute to complex trait variation. The aggregate effect of these regions, however, may only explain a small fraction of the expected heritable variation, a phenomenon referred to as “missing heritability” [1]. The potential sources of the missing heritability are varied and include; rare variants, structural variation, epistasis, and the focus of this article, epigenomics [1–3].

Recently, there has been great excitement investigating the role of possible epigenomic sources of variation, in the form of stable DNA or histone modifications, which may act alongside, or independently, of traditional genetic variation. A range of population-level studies have reported substantial diversity between different genetic backgrounds [4–9]. In certain cases, measures of DNA methylation were combined with genetic variation identifying genetic factors that can influence chromatin between populations and geographical locations [6,10–12]. Results often highlight the covariation of genetic variation with chromatin state, transposable element (TE) methylation, and differential cytosine methylation among accessions. Nonetheless, a small portion of genetically-independent methylation variation, often in the CG context and in promoter regions, may contribute to phenotypic variance [6,13–15].

To date, the major advances in the field of epigenomics have rarely shown a direct relationship with downstream phenotypes [16,17]. In addition, changes in chromatin state are often found to be confounded with genetic variation present between samples. That is, chromatin variation is clearly apparent in cases where genetic variation exists and it has been difficult to disentangle these sources of variation when associated with a phenotype of interest [18]. Many studies address this by examining epigenomic variation through inbreeding generations, so called epi-Recombinant Inbred Lines (epiRILs) or mutation accumulation lines, which can display phenotypic variation [13,19–21]. However, such estimates of epigenomic variation on phenotypic differences are often made in isolation. Given the current state of epigenomic research, a more holistic approach in which chromatin variation is assessed along with genetic polymorphisms and environmental variation to reveal the relative importance of these heritable factors. The ‘Extended Genotype’ of an organism can be defined as sources of heritable variation that are largely overlooked and/or misinterpreted in relationship to more traditional genotype assessments such as SNPs [18].

The model grass *Brachypodium distachyon* (*B. distachyon*) provides an ideal system to examine the impact of the extended genotype on natural populations. Advantages include having a relatively small and fully sequenced genome alongside a growing array of genetic resources, displaying a wide climatic distribution resulting in phenotypic diversity, rapid generation times leading to increased rounds for (epi)genetic selection, and a small stature that facilitates systematic study. Pertinently, multiple genetically similar accessions of *Brachypodium* have been identified in different environments across Turkey [22,23]. This set of ‘BdTR’ accessions provide a unique natural set of germplasm to investigate the impact of ‘extended genotype’ signals, such as DNA methylation.

## Results

### Selection of *Brachypodium* accessions and genetic profiling

An initial genomic analysis of thousands of *Brachypodium* accessions was conducted using genotype-by-sequencing (GBS) culminating in a selection of diverse *B. distachyon* germplasm as a *HapMap* set [22]. In that study, it was observed that some Turkish accessions were highly similar based on SNP assessment, but were located in different geographical regions of Turkey (Figure 1A). These 83 Turkish accessions (BdTR) were grouped into seven nearly genetically identical ‘families’ based on prior analysis (Sup Table 1). This was generally consistent with the accession’s initial naming based on phenotypic similarity [23], though BdTR1 and BdTR2 belonged to the same genotype family. The BdTR accessions within each genetic family were widely dispersed geographically and found at different elevations (Figure 1B) consistent with wide migration and little recombination of the highly selfing species.

**Figure 1:**
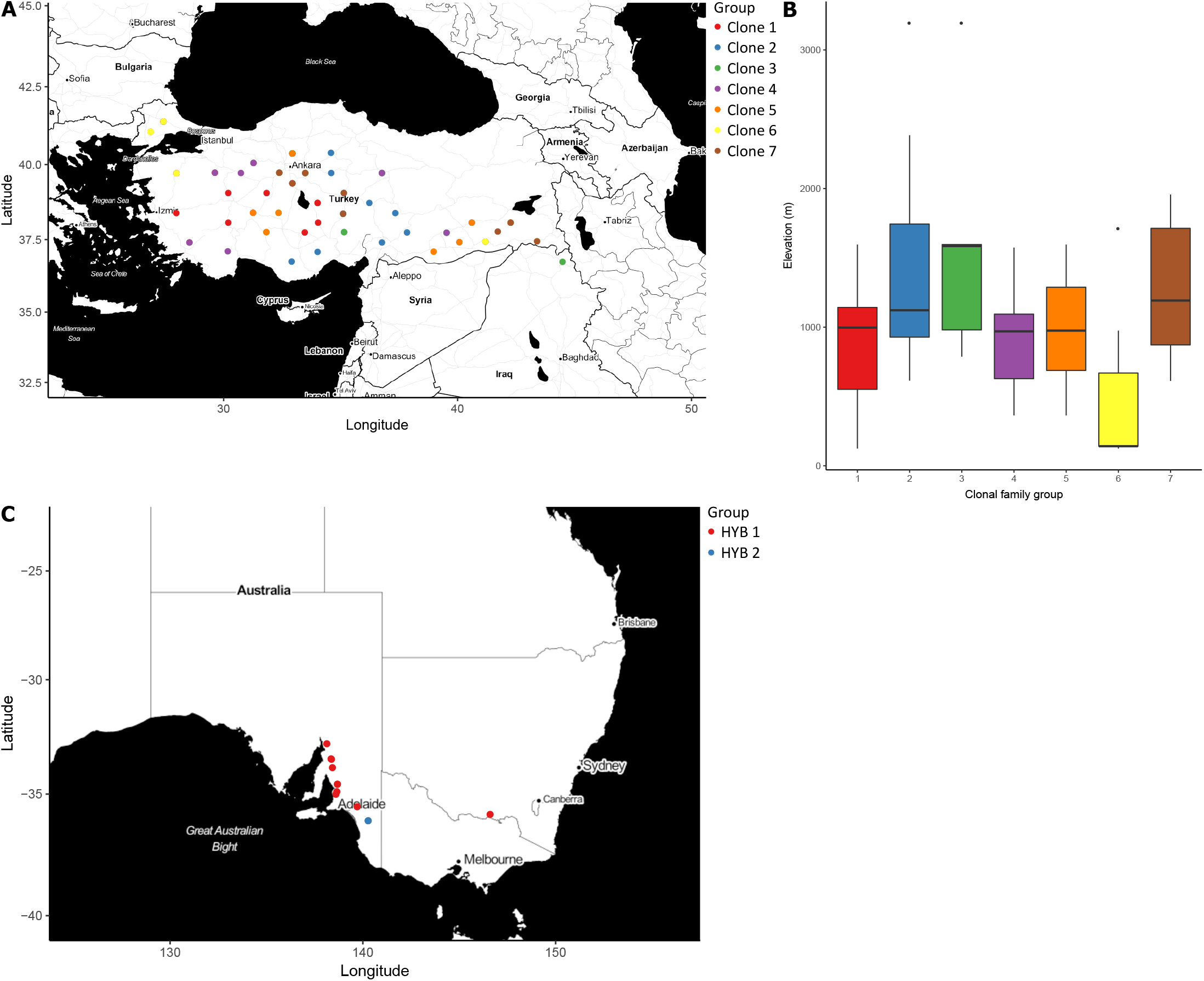
Selection of *Brachypodium* accessions for analysis. (A) Map of diploid Turkish accessions colored by genetic relationship. (B) Boxplot of genetic family elevation variation from initial collection sites. (C) Map of *Brachypodium hybridum* accession collection locations. Colors indicate genetic grouping into ‘H1’ or ‘H2’ subsets.

As a genetically distinct global outgroup, we included the allopolyploid relative *Brachypodium hybridum* (*B. hybridum*). Previous studies highlighted two genetically-similar families consisting of 12 accessions collected from across southern Australia. HYB1 consisted of accessions initially collected in South Australia and one location in New South Wales. HYB2 consisted of a single geographic location with multiple individuals (Figure 1C). The selection of germplasm for this study provided a unique, natural system in which to study new heritable variation across a series of genetic families containing minimal *intra-family* variation within, but with substantial *inter-family* genetic and phenotypic variation.

### Methylome variance reflects genetic distance

To examine the DNA methylation state of accessions, low-coverage (~1-2x) whole genome bisulfite sequencing was conducted for all 604 experimental plants (Sup Table 1). Samples were correlated over 100 bp genomic tiles to obtain average genome-wide methylation state for all three sequence contexts. CG methylation almost completely reproduced the known genetic relationships - separating B. hybridum and grouping the B. distachyon accessions based on previously determined family clone groups (Figure 2A). Individual biological replicates were often most similar, however, variation observed near terminal edges may be due to the limitations of low coverage sequencing data. This is apparent in the non-CG methylation contexts (CHG and CHH) in which sample relationships showed less complete recapitulation of genetic relatedness (Figure 2B-C). This was likely due to low coverage preventing accurate measurements for these methylation contexts that occur at lower levels. Therefore we focus the remainder of the analysis on CG sites.

**Figure 2:**
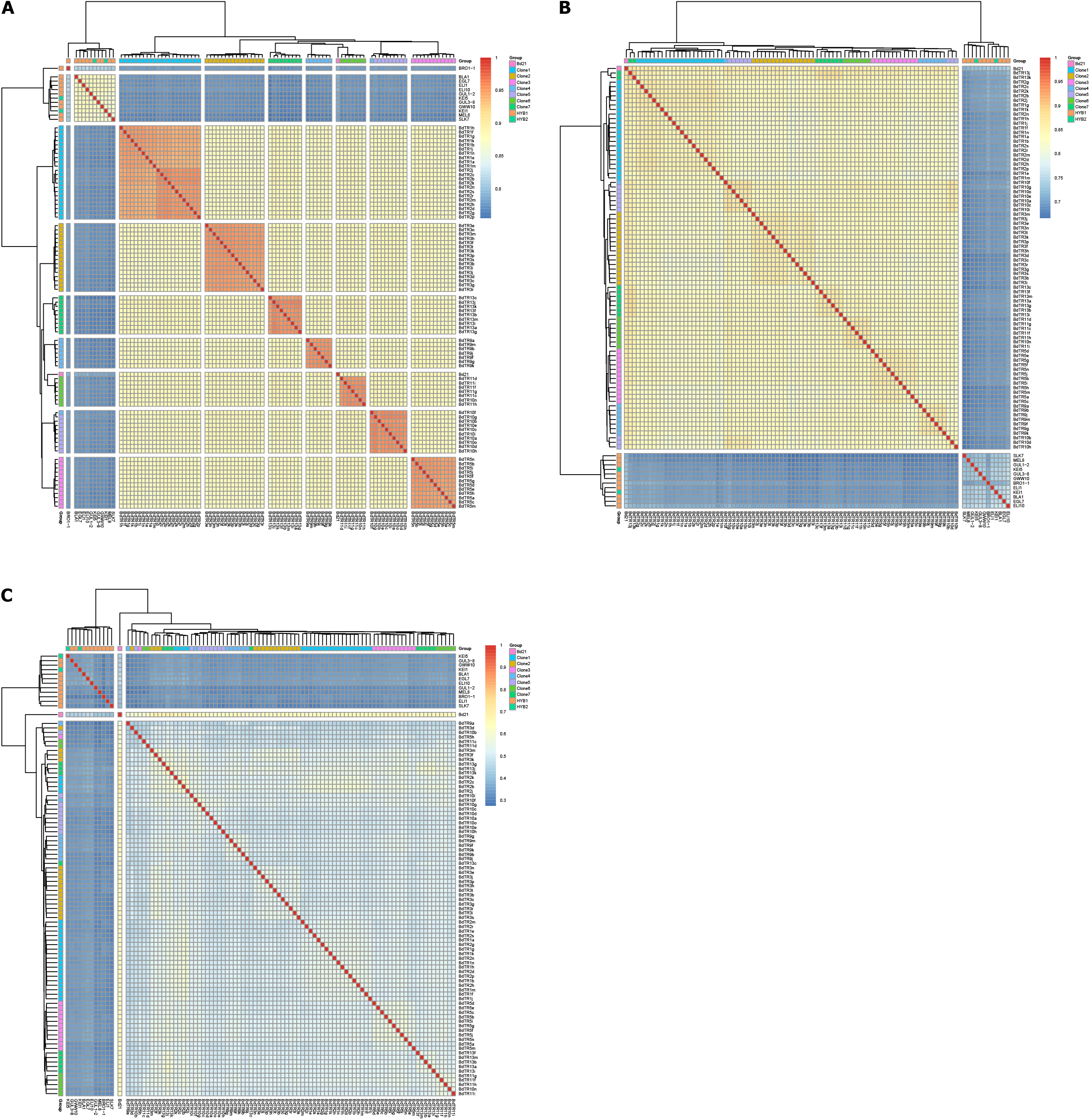
Broad methylation patterns resemble genetic relationships. Heat maps representing two-dimensional hierarchical clustering of *Brachypodium* accessions based on correlations (Pearson’s r) in genome-wide DNA methylation levels, binned into 100 bp genomic tiles and averaged across all replicates per accession, for (A) CG, (B) CHG, and (C) CHH contexts. Family groups are denoted based on known genetic relationships.

The nested design of this experiment allowed for further quantification of DNA methylation variation at a variety of levels. We identified differentially methylated regions in the CG context (CG-DMRs) using single-sample *HOME* DMR calling for all pairwise comparisons between: (I) replicates of the same accession (basal level of stochastic differences), (II) accessions within families (*intra*-family), and (III) between families (*inter*-family, Table 1, Figure 3). This was performed for the complete dataset (low coverage) and repeated for a subset of samples (family group 1 and 6) sequenced to greater depth (~10X). Indeed, greater sequencing depth improved the power of *HOME* to detect CG-DMRs. As comparisons were made between samples with increasing genetic distance, a greater mean number of DMRs were identified (of greater magnitude, length, and CG count). On average, 4-fold more CG-DMRs were called between, compared to within, family groups, highlighting previous observations that genetically variable samples contain many more DNA methylation variants. Nonetheless, a substantial number of CG-DMRs were still observed within accessions and family groups, which could contribute to heritable and even adaptive phenotypic variation. This warranted further systematic analysis to quantify the phenotypic variance explained by genetic and epigenomic components.

**Figure 3:**
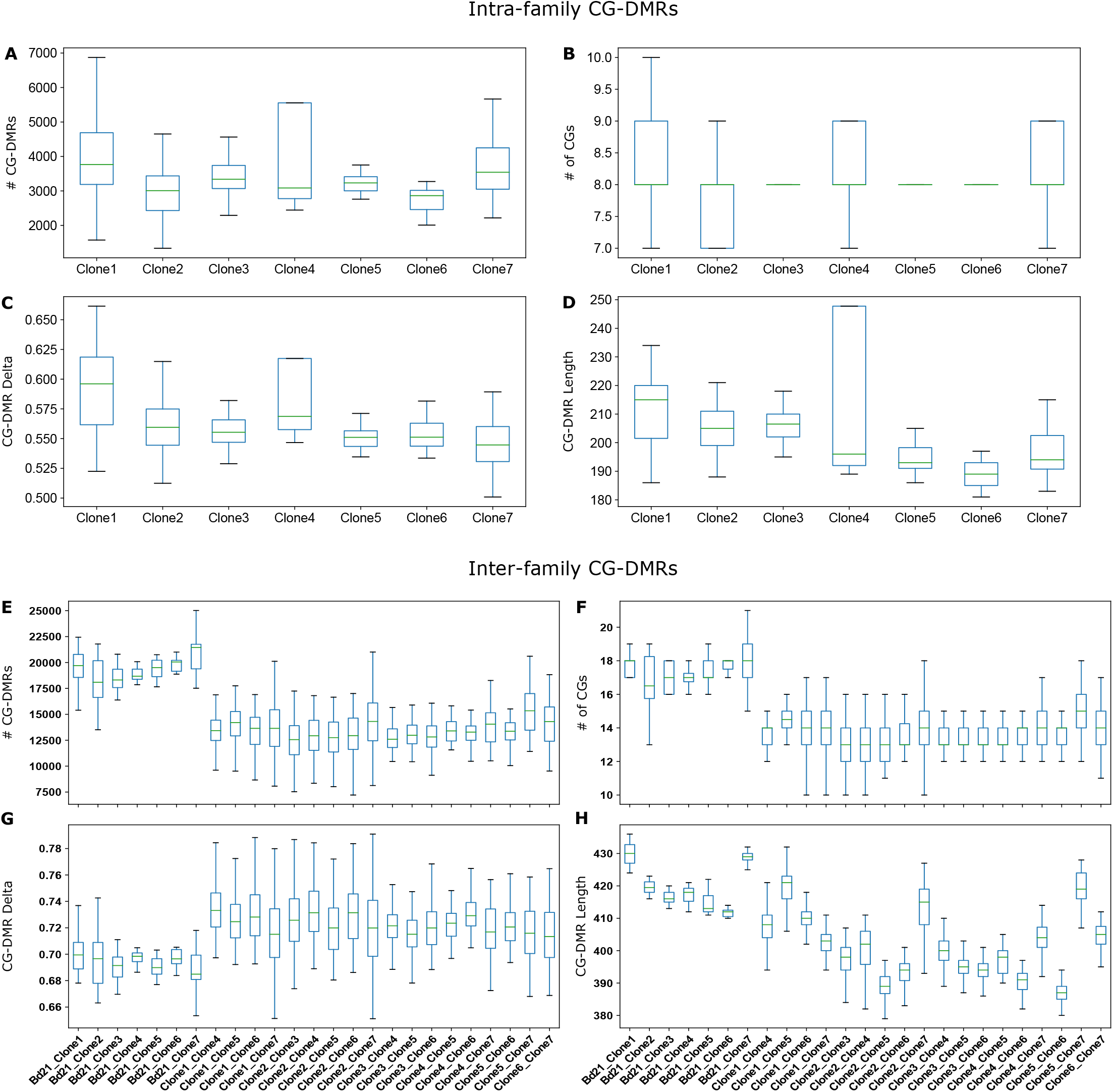
Pairwise intra- and inter-family CG-DMRs. Box plots presenting the results of pairwise CG-DMR calling with HOME for intra- (A-D) and inter-family (E-H) comparisons. Plots display the distribution of the number of CG-DMRs (A, E), number of CGs per CG-DMR (B, F), and delta (C, G) and length (D, H) of CG-DMRs.

**Table 1.**
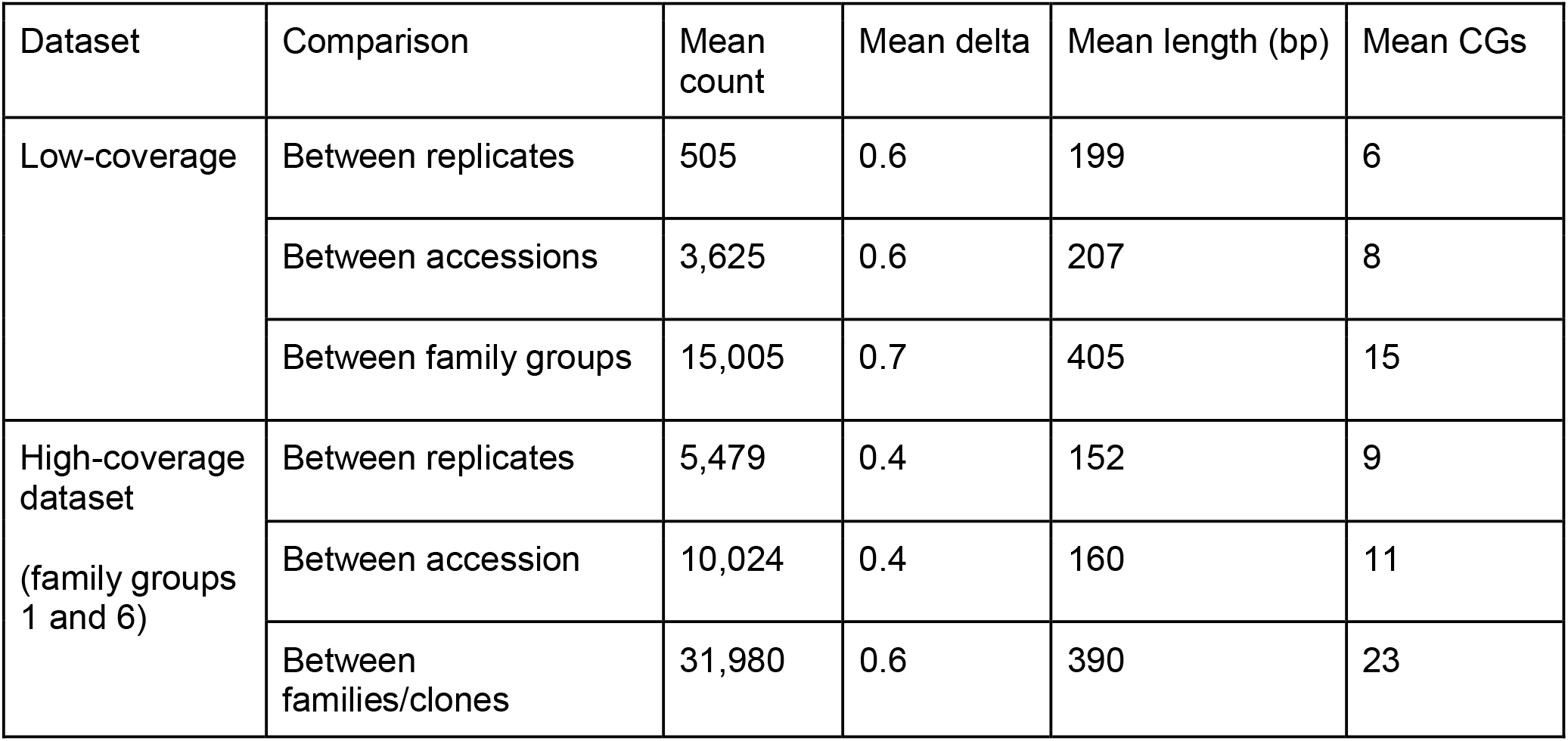
CG-DMR analysis using *HOME*

### Dissection of phenotypic variance into genetic and epigenomic components

We used a mixed linear model framework to dissect the phenotypic variance of fitness related traits across multiple levels of biological organisation. Our experimental design allowed us to investigate the differences between the species *B. distachyon* and *B. hybridum*, and within our *B. distachyon* samples. We estimated the relative contribution to phenotypic variance of additive genetic effects (as characterised by whole-genome sequence distance among seven clone groups), CG methylation state, which was obtained through low-coverage bisulfite sequencing of 83 accessions, and the residual variance being estimated via the biological replicates of the accessions. In the analyses that considered all hierarchical levels of variation, the species contrast of *B. distachyon* and *B. hybridum* explained most of the phenotypic variance in flowering time in spring conditions (84%) where *B. distachyon* flowered considerably later than *B. hybridum*. A similar pattern was observed in the fall overwintering conditions; however, this relationship explained less variation overall (20%) as vernalization overwhelms the genetic signal. We also found differences in leaf morphology between the two species, where *B. hybridum* had both wider and longer leaves compared to *B. distachyon*. This difference in leaf width and length between the species accounted for 68% and 94% of variance in spring conditions and 76% and 98% in fall conditions for these traits respectively.

Within *B. distachyon*, we observed substantial genetic variance across all traits in both conditions, where heritabilities ranged from 18% to 63% in spring conditions and from 10% to 89% in fall conditions (Table 2). The addition of the random effect of CG methylation state to the models rarely increased the amount of phenotypic variance explained compared to the simpler model where only additive genetic effects were considered. In this study, only two traits had statistical support that the phenotypic variance explained by methylation patterns was greater than zero and, in both cases, this result was dependent on the conditions in which the traits were measured (Table 2). We estimated that an additional 10% of variation in flowering time is due to methylation in spring conditions (◻^2^ = 7.29, d.f. = 1, p = 0.007), we also found methylation patterns explained an additional 16% of the variation in plant height measured in fall conditions (◻^2^ = 13.14, d.f. = 1, p < 0.001). We would like to stress the importance of the joint estimation of genetic and methylation contributions to phenotypic variance as these two explanatory variables were highly correlated, i.e. closely related individuals shared similar methylation patterns. In this study, ignoring the underlying DNA sequence variation in our sample population would have led us to grossly overstate the importance of methylation state on phenotypic variance. For example, we found that methylation state only explained 5% of variation in the tiller count in fall conditions (◻^2^ = 3.39, d.f. = 1, p = 0.065), while additive genetic effects explained 60% of the variation (Table 2). Re-analysing the data with the removal of the random effect of additive genetics from the model, the amount of variation explained by the methylation state dramatically increased to 51% (◻^2^ = 133.33, d.f. = 1, p < 0.001). Overall, we show that in some cases significant phenotypic variation can arise due to epigenomic processes that can potentially be inherited by subsequent generations, however, we stress that variation in methylation state is largely dependent on the underlying genetic variation and needs to be analysed together.

**Table 2.** Sources of phenotypic variation. Breakdown of the phenotypic variance from the independent effect of SNPs (i.e broad-sense heritability), the equivalent effect of CG methylation state, and environment variance. Variance components were estimated using linear mixed models and have been mean-standardised and multiplied by 100 for readability. Bold components were found to be statistically significant at P < 0.05 using log-likelihood ratio tests between full and reduced models.

| Trait | Spring | Fall |
| --- | --- | --- |

| | $I_A$<br>($H^2_{\text{SNP}}$) | $I_M$<br>($H^2_{\text{CG}}$) | $I_{\text{RES}}$ | $I_A$<br>($H^2_{\text{SNP}}$) | $I_M$<br>($H^2_{\text{CG}}$) | $I_{\text{RES}}$ |
| --- | --- | --- | --- | --- | --- | --- |
| Flowering time | **0.332**<br>**(0.485)** | **0.062**<br>**(0.096)** | 0.269 | **0.245**<br>**(0.894)** | 0.004<br>(0.013) | 0.026 |
| Height | **1.384**<br>**(0.387)** | 0.124<br>(0.035) | 2.073 | **0.327**<br>**(0.373)** | **0.140**<br>**(0.161)** | 0.407 |
| Tiller count | **3.984**<br>**(0.633)** | 0.311<br>(0.056) | 1.992 | **4.240**<br>**(0.597)** | 0.365<br>(0.051) | 2.494 |
| Leaf length | **0.572**<br>**(0.483)** | 0.048<br>(0.041) | 0.563 | **0.538**<br>**(0.523)** | 0.055<br>(0.053) | 0.437 |
| Leaf width | **1.736**<br>**(0.627)** | <0.001<br>(<0.001) | 1.035 | **1.015**<br>**(0.241)** | 0.379<br>(0.090) | 2.818 |
| Ear count | **0.806**<br>**(0.176)** | 0.308<br>(0.067) | 3.474 | **0.290**<br>**(0.100)** | 0.232<br>(0.086) | 2.273 |

## Discussion

Epigenomic diversity continues to be considered as a new source of variation in heritable traits that could be harnessed for plant breeding [24]. However, genetic and epigenomic polymorphisms are often considered independently making it difficult to determine their relative contribution towards heritable phenotypic variation. Here, utilizing a diverse range of clonal *Brachypodium* accessions, grown in two distinct controlled environments (simulating local climates of Istanbul spring and fall), we systematically quantified the proportion of phenotypic variance, across numerous adaptive plant traits, attributable to genetic polymorphisms (SNPs), epigenomic variation (CG methylation), and the environment. Whereas the majority of phenotypic differences across all traits could be attributed to genetic polymorphisms, CG methylation demonstrated an additive effect in particular environments, such as plant height under longer overwintering (fall) conditions.

Although the known population structure of the selected accessions in this study was recapitulated with methylation data, the level of methylation variation was linked to the overall genetic distance between any given samples. This result is similar to reports of natural *Arabidopsis thaliana* populations [6,12,25] and across a landrace Wheat collection [26]. Methods as described here, can dissect this confounding between genetics and epigenomics, to capture an additional piece of the missing heritability. This highlights the importance of dissecting chromatin variation along with correlated genetic variation to explain phenotypic variation. In the majority of cases where samples are genetically similar, phenotypic variation of quantitative traits is also limited. Despite this, our analyses also revealed that epigenomic distance (CG methylation) could explain additional phenotypic variation of particular traits in particular environments.

Going forward, in experimental systems in which genetic sources of variation are not known, it would be advantageous to separate epigenomic (base modifications) from genomic changes (SNPs, SVs) to be able to jointly test effects on a phenotype. Fortunately, advances in genomics, including long reads that also type base modifications, are upon us making this a practical solution [27–29]. In essence, a joint understanding of genetic and epigenomic relationships could be a new standard for the examination of quantitative trait variation.

## Methods

### Plant germplasm, growth, and phenotyping

Germplasm for experiments was selected based on previously established SNP-based genotypic relationships as described in [22]. Briefly, the genetic distance matrix, as defined for all samples, was used to identify clades of individuals with genetic variation similar to that of technical replicates and for which seed was available. The 95 selected *B. distachyon* accessions were grown in triplicate alongside 17 biological replicates of the reference genotype Bd21 for direct comparison to an inbred background [31]. Two seeds per pot were planted ~2.5 cm below surface in 5×5×8 cm pots in a steam-pasteurised 75:25 martin’s soil:washed river sand mix. Trays of 14-16 pots were watered with tap water, covered in plastic film, and placed at 4 °C in the dark for seven days for seed stratification. Trays were subsequently moved into modified Conviron growth chambers which have been fitted with 7-band LED light panels and control light intensity, quality, chamber temperature, and humidity every 5 minutes [32]. These were planted under simulated conditions for regions near Istanbul, Turkey season-shifted for planting in both northern-hemisphere spring (April) and fall (August) to investigate how plants respond to either a rapid-cycling or overwintering environment. A pair of identical specialized growth chambers were used to simulate these Turkish climates (41.146, 29.026, 72m elevation) modeled using SolarCalc (version E-2014 [33]). The chambers updated temperature and 7-band LED light quality information every 5 min (Sup Figure 1) [32]. Spring started with average high temps of 17 °C and reached a peak summer daily high at ~29 °C. Night time lows ranged from a season start of ~10 °C to ~21 °C by the peak of summer (simulated July). In contrast, fall planting started with average high temps of 19 °C and reached a minimum winter daily high at ~7 °C. Night lows ranged from a season start of 14 °C to 5 °C (limited by chamber specifications) by mid-winter (simulated February). The two chambers performed the same modeled conditions offset by six months. This allowed for a ‘spring’ and ‘fall’ planting in which plants entered the simulated conditions on April 26th and October 16th. Plants were regularly watered using tap water over the course of the experiment when standing water was not observed in trays. All plant measurements were taken by hand, including plant height, third leaf length and width, tiller count, ear count, and flowering time (Sup Table 2) [22,34,35].

### Tissue Harvest and DNA extraction

The flag leaf or leaves 3 and 7 were harvested for spring and fall plantings, respectively. Tissue was harvested directly into 96 well plates on liquid nitrogen for DNA extraction. Leaf tissue was ground using the TissueLyser II (Qiagen) and DNA was extracted using the Invisorb DNA Plant HTS 96 Kit (cat 7037300400; Stratec Biomedical) using the manufacturer’s protocol. DNA was quantified using the Quant-iT High Sensitivity dsDNA Assay (cat. Q33120; ThermoFisher).

### Whole genome bisulfite sequencing and analysis

Whole genome bisulfite sequencing libraries were created using the Accel-NGS Methyl-Seq DNA Library Kit (cat. 30096; Swift Biosciences; Ann Arbor, MI). The standard protocol was modified for third-reaction volumes throughout using 27 ng bisulfite converted gDNA. Initial shearing step of gDNA was omitted. Bisulfite conversion was conducted using the EZ-96 DNA Methylation-Gold MagPrep kit (cat. D5043; Zymo Research). 80 ng of gDNA in 45 μl H_2_O was used for low concentration gDNA. Dual indexing of samples was performed with the Methyl-Seq Dual Indexing Kit (cat. 38096; Swift Bioscience) using 11 PCR cycles. The library underwent a final dual-side SPRI cleanup upon completion of the library preparation (0.6x right-side SPRI followed with 0.85x left-side SPRI) to compensate for the lack of physical shearing of the initial DNA. Libraries were quantified using the Caliper LabChip GXII (PerkinElmer) and equal-molar pooling of 96 libraries was performed. Pools of 96 libraries were sequenced using the HiSeq 2500 (Illumina). The subset of samples selected for high-coverage methylome analysis were run as a pool of 96 samples across a flow cell (8 lanes) on the HiSeq 2000. All sequencing was conducted at the ANU Biomolecular Resource Facility.

Raw sequencing reads had 5’ trimming of 6 bp to eliminate library bias in methylation state, along with base quality and adapter trimming using *Trim Galore!*. Trimmed reads were mapped to the Bd21 v2.1 using Bismark (v0.13.0) under bowtie1 alignment mode [36]. Alignments were subsequently deduplicated and all context (CG/CHG/CHH) methylation data was extracted via bismark_methylation_extractor using default parameters.

DMRs were called using HOME-pairwise module with --delta 0.2 --minc 4 and --len 50 [37]. HOME is a machine learning based DMR identification method which accounts for biological variation present between the replicates and uneven read coverage through weighted logistic regression while computing the P-value. The spatial correlation present among neighboring cytosine sites is captured by moving average smoothing and the use of weighted voting for histogram based features [37].

### Dissection of phenotypic variance

We used linear mixed modeling to estimate the amount of phenotypic variance that can be explained by the differences in CG methylation between accessions while accounting for the genetic effects due to differences between the clone groups. Analyses were performed for each trait and environment separately and was fitted using the R package “*Asreml-R*”. Initially, we construct a reduced model with one random effect:

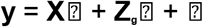

Where **y** is a vector of phenotypes, ⍰ the population mean, ⍰ is the vector of breeding values treated as a random effect with a ~N(0, σ^2^**G**) distribution, and ⍰ is the vector of residual effects. **Z**_**g**_ is the design matrix allocating clone groups to individual plants and has been defined as the inverse of the genomic relatedness matrix **G**. The genomic relatedness matrix was estimated following [38] that has been adjusted for almost completely homozygous organisms. First, from whole-genome sequencing data from one accession of each clone group, we build the matrix **X** with the dimensions of number of clone groups (n=7) and number of SNPs (m=510,230). SNPs with at least one copy of the minor allele and no more than one missing value were included in **X**. **X** was rescaled to account for allele frequencies to produce **W** where W_ij_ = M_ij_ − p_j_, where p_j_ is the allele frequency for SNP j. Finally, **G** is calculated as **G** = **WW’**/∑p_j_(1-p_j_).

We then construct a more complicated model with an added random effect to capture the variance due to differences between the accessions in CG methylation states:

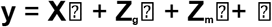

Here, ⍰ is a vector of fitted values (~N(0, σ^2^**M**) that we interpret as the methylation equivalent of breeding values for which the “methylation heritability” can be estimated from the variance in these values. **Z**_**m**_ is the design matrix allocating accessions to plants defined by the inverse of a methylation similarity matrix described below.

We pooled the reads from bisulfite DNA sequencing of the replicates of each accession separately for each environment. For each accession, we scored the proportion of methylated reads compared to the total number of reads at all CG sites. We further pooled CG sites into 200bp windows. The proportion of methylated reads for a specific window was treated as missing data if an accession had less than 10 reads total. Windows with more than 50% of accessions containing missing values were excluded from further analysis. Additionally, windows that did not contain a single methylated read were also excluded. In total, 838,231 windows were retained in the spring conditions and 778,183 windows in the fall conditions.

After filtering, we retained a matrix of proportions of methylated reads, **Q**, with the dimensions of accessions (n=83) by the number of 200bp windows. Columns of **Q** were scaled to have a mean of zero and unity of variance. The methylation similarity matrix was then calculated as **M** = **QQ’**/**N**, where **N** is a matrix of pairwise number of windows with non-missing data between two accessions.

We used a log likelihood ratio test to determine if the methylation state of the accessions explained a statistically significant amount of phenotypic variance. Here the statistic of 2 times the difference in log likelihood between the models was tested against a chi-squared distribution with one degree of freedom. All variance components were mean-standardised following [39].

### Data Accessibility

All bisulfite sequencing data has been deposited at the NCBI under BioProject PRJNA349755. All phenotypic and experimental records can be found at github.com/steichten/clonal_brachypodium.

## Acknowledgements

The authors would like to thank the Australian Plant Phenomics Facility for providing plant growth facilities and the National Computational Infrastructure for provision of computing resources, both of which are supported under the National Collaborative Research Infrastructure Strategy of the Australian Government. Illumina sequencing was performed at the Biomolecular Research Facility at the ANU. SRE was funded by an Australian Research Council Discovery Early Career Research Award (DE150101206). This project was supported by the Australian Research Council Centre of Excellence in Plant Energy Biology (CE140100008).

The authors declare no conflicts of interest.

## Figure Legends

**Supplementary Figure 1:**
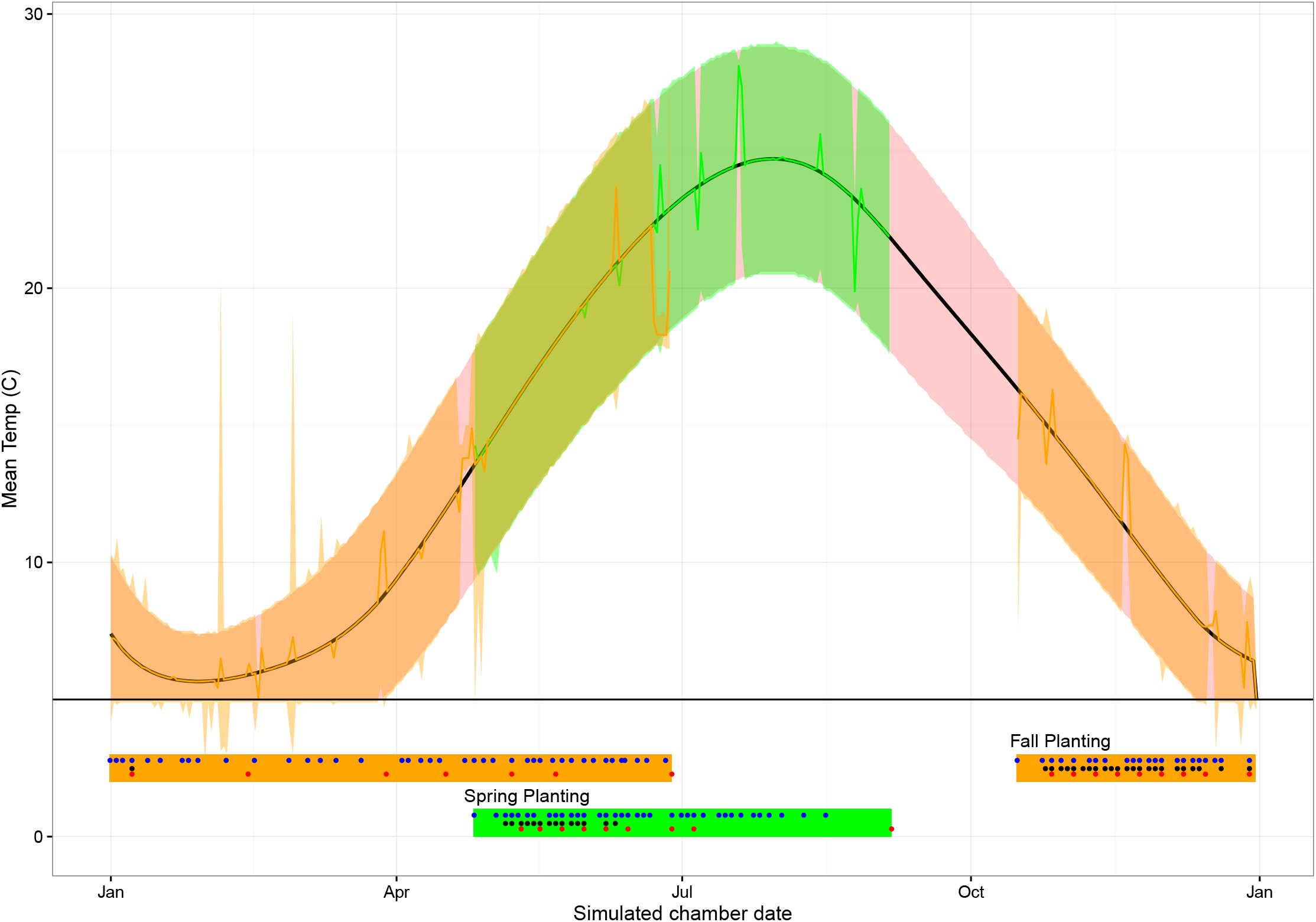
Growth chamber conditions and planting summary. Curve indicating the overall temperature settings through a year of simulated Turkish conditions. Black curve indicates per-day temperature averages with red boundaries indicating set minimum and maximum temperatures within a 24-hour cycle. Orange and green bands highlight the simulated fall and spring planting and growth period chamber temperatures respectively. Deviations outside of the red boundaries highlight technical limitations or errors in the chamber temperature program. Horizontal line at 5°C is the lowest approved temperature setting of the chambers. Lower panel highlights fall and spring growing periods. Dots indicate dates where watering (blue), plant height (red), and growth stage (black) were measured.

